# Circuit-free, optoelectronic sensing of electrodermal activities based on photon recycling of a micro-LED

**DOI:** 10.1101/2020.07.05.188391

**Authors:** He Ding, Guoqing Lv, Zhao Shi, Dali Cheng, Yunxiang Huang, Lan Yin, Jian Yang, YongTian Wang, Xing Sheng

**Affiliations:** Beijing Engineering Research Center of Mixed Reality and Advanced Display, School of Optics and Photonics, Beijing Institute of Technology, Beijing 100081, China; Department of Electronic Engineering and Beijing National Research Center for Information Science and Technology, Tsinghua University, Beijing 100084, China; School of Materials Science and Engineering, Tsinghua University, Beijing 100084, China

**Keywords:** optoelectronic sensing, photon-recycling, micro-LEDs, galvanic skin response

## Abstract

Conventional epidermal electronics integrate multiple power harvesting, signal amplification and data transmission components for wireless biophysical and biochemical signal detection. This paper reports the real-time electrodermal activities can be optically captured using a microscale light-emitting diode (micro-LED), eliminating the need for complicated sensing circuit. Owing to its strong photon-recycling effects, the micro-LED’s photoluminescence (PL) emission exhibits a superlinear dependence on the external resistance. Taking advantage of this unique mechanism, the galvanic skin response (GSR) of a human subject is optically monitored, and it demonstrates that such an optoelectronic sensing technique outperforms a traditional tethered, electrically based GSR sensing circuit, in terms of its footprint, accuracy and sensitivity. This presented optoelectronic sensing approach could establish promising routes to advanced biological sensors.

---

Epidermal sensing systems based on advanced electronic and photonic materials and devices have gained significant interest in the past decade, owing to their versatile capabilities in monitoring biophysical (electrophysiology, temperature, pressure, etc.) and biochemical (oxygen, glucose, ions, etc.) activities with direct clinical relevance.^[1-4]^ To realize remote, non-invasive, precise and continuous operation, various strategies utilizing microwave, optical, and ultrasound signals have been developed for wireless energy harvesting and signal transmission ^[5-9]^. However, conventional skin-mounted sensors, or bioelectronic sensors in general, still require sophisticated circuits with multiple device components, including at least a power supply, a signal amplifier and a data transmitter.^[10]^ Alternatively, it is known that a semiconductor photodiode can serve as an energy generator (via the photovoltaic effect) and a light emitter (via the radiative recombination of carriers).^[11]^ Moreover, for a diode with high electron-to-photon and photon-to-electron conversion efficiencies, its photoluminescence (PL) can be leveraged by manipulating its photon-recycling effects based on varied electrical signals like voltage, current and resistance.^[12-16]^ In other words, this implied that such a mechanism could provide an unusual way for probing electrophysiological signals.

In this paper, we demonstrate that the electrodermal activity (skin resistance, or galvanic skin response GSR) can be optically monitored by simply using a thin-film, microscale light-emitting diode (micro-LED), with a footprint of about 100 micrometers (**Figure 1**a). Such a micro-LED can simultaneously work as an energy absorber, a signal amplifier and an optical transmitter. The resistance dependent photon-recycling effect alters the micro-LED’s luminescence under optical excitation. With its electrodes attached to the human skin, its optical signals can directly reflect the real-time GSR variations, with performance comparable to a commercial GSR sensing circuit module. This concept would provide a novel optical solution to circuit-free, wireless biosensors.

**Figure 1.**
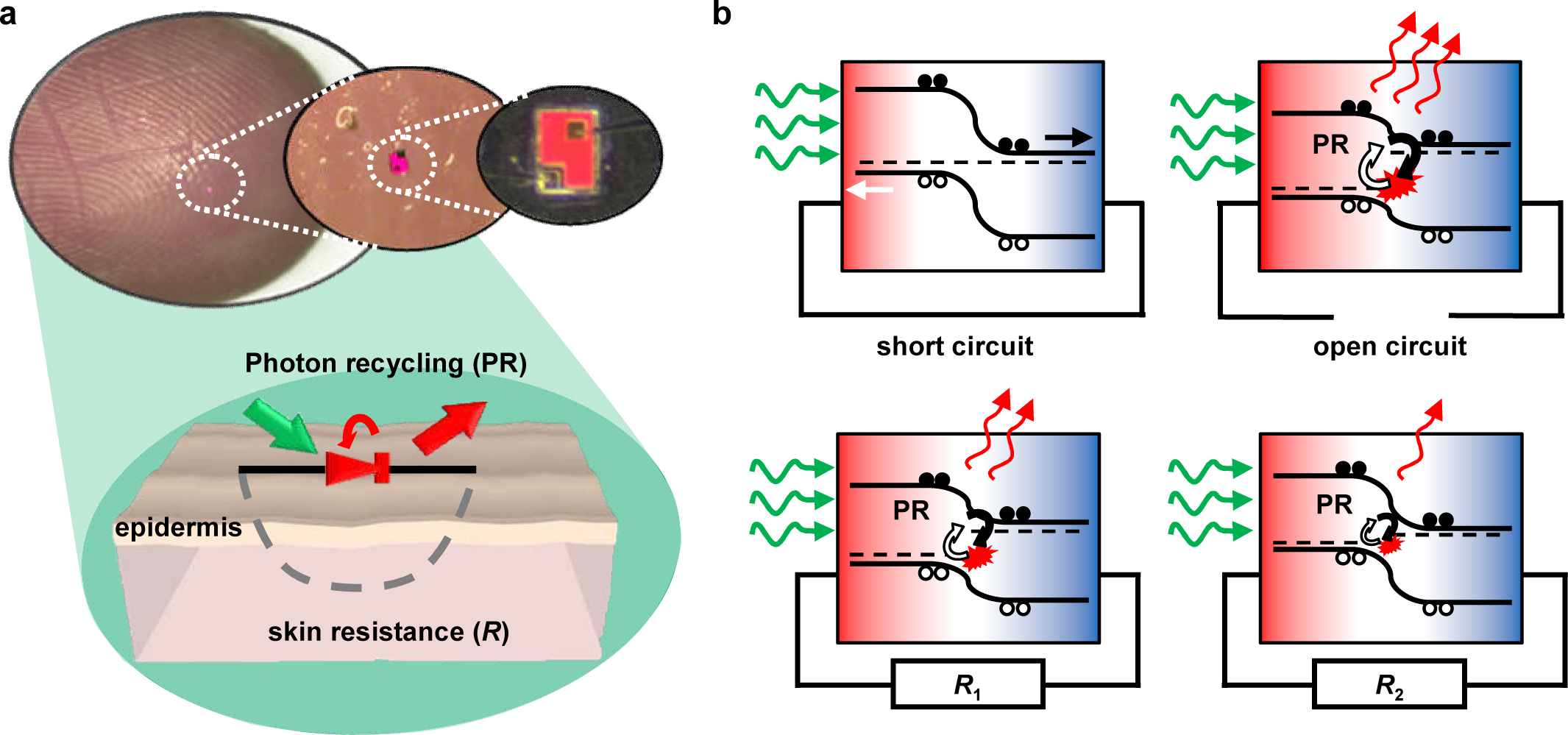
(a) Images and schematic illustrations of a thin-film, red-emitting micro-LED attached onto the human skin for wireless, optoelectronic sensing of the galvanic skin response (GSR), based on its resistive dependent photon-recycling (PR) effect. The LED has lateral dimensions of 200 μm × 150 μm and a thickness of 5.6 μm. (b) The operational principle of the sensor, showing that the diode’s photoluminescence (PL) intensity is dependent on its load resistances in different working conditions: short-circuit, open-circuit, and connected to different resistances (*R*_1_ > *R*_2_). Black and white dots indicate photogenerated free electrons and holes, and red, blue and white regions represent the p-type, n-type and depletion regions of the diode, respectively.

Figure 1a shows photographs of the gallium indium phosphide (GaInP) based red-emitting micro-LED attached on the human finger. The thin-film, freestanding GaInP micro-LED has a lateral dimension of ∼200×150 μm^2^ and a thickness of ∼5.6 μm, fabricated by epitaxial growth, lithographic etching, metal deposition, removal of the sacrificial layer and transfer printing (more details see the methods).^[17]^ For circuit-free operation, no current injection is provided through external power supplies like a battery. Instead, the micro-LED is optically illuminated with a green laser source, and its red PL emission can be captured through a 560 nm long-pass filter.

Schematically illustrated in Figure 1b, its operational principle as a wireless GSR sensor can be understood based on the modeling of a simplified photodiode under optical excitation. In the short-circuit condition, most of the photogenerated carriers flow into the external circuit and recombine non-radiatively. At the open-circuit condition, on the contrary, the carriers recombine within the junction via photon-recycling, creating a strong PL if the diode is made by III–V semiconductors with high external luminescence efficiencies (*η*_ext_).^[18-20]^ In other words, the load resistance *R* alters the diode’s build-in potential, thereby modulating its PL emission (PL intensity increases with *R*). In detail, the relationship between the diode’s PL intensity and the loaded skin resistance *R* can be described by (see details in the Supplement information 1):

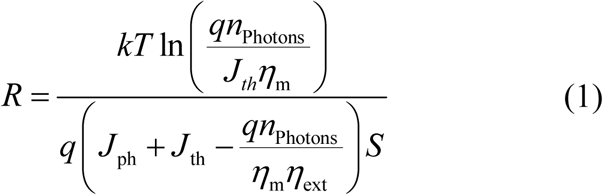

where *J*_ph_ is the photogenerated current derived from the excitation light power density, *J*_th_ is the absorbed thermal radiation from the environment, *S* is the active surface area of the device, *n*_photon_ is the number of the captured emissive photons directly related to the PL intensity and the coupling efficiency of the microscope setup (*η*_m_). It can be seen that an exponential relation between the PL emission and the resistance is obtained due to the diode characteristics, providing an effective amplification mechanism for GSR sensing.

**Figures 2**a and b present the details about the GaInP micro-LED structure, with more information in the Table S1. We evaluate its GSR sensing capability by connecting it with different load resistances (Figure 2c). Under the green light excitation (peaked at ∼ 545 nm) with fixed power, the micro-LED’s PL emission increases with the increased resistance values and reaches the maximum at the open-circuit condition (Figure 2d). We further measure the integrated emission intensity (unit: counts/μm^2^/ms) as a function of the resistance at various excitation power, using a fluorescence microscope equipped with a CMOS camera (details see in the methods). The results are plotted in Figure 2e, in comparison with the calculations based on Equation 1. The measured PL intensity firstly increases exponentially with the resistance and then saturates when approaching the open-circuit, which is in accordance with the theoretical predictions. In addition, there is a trade-off between the resistance sensitivity and the detection limit of the measurement setup (∼5–10 counts/μm^2^/ms). As shown in the literatures^[21, 22]^ and subsequent experiments, the GSR levels for the human body normally lie within 100–500 kΩ; therefore, an excitation power of ∼1.91 mW/mm^2^ is selected for actual GSR measurement involving human subjects.

**Figure 2.**
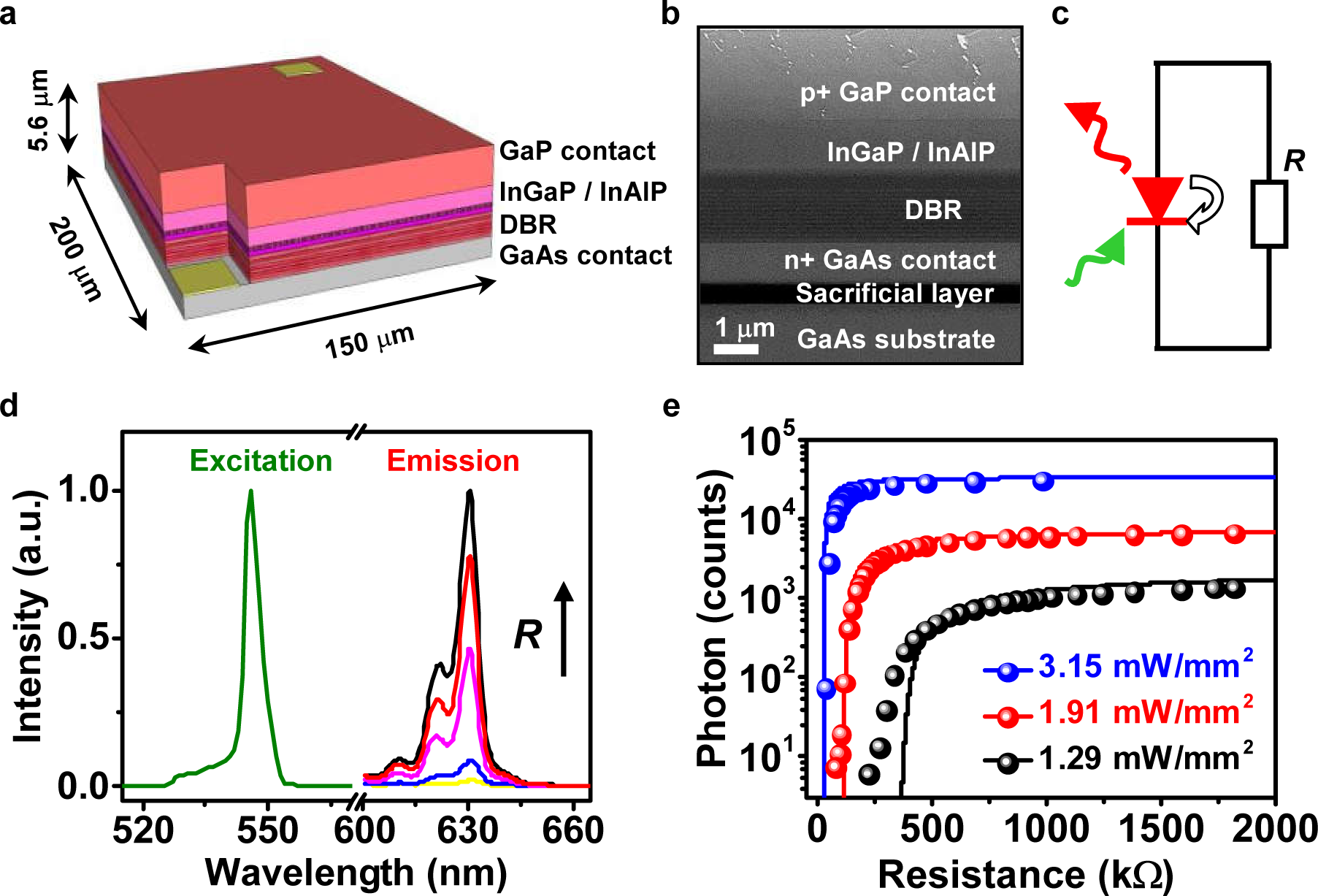
(a) Schematic diagram and (b) cross-sectional scanning electron microscope (SEM) image of the micro-LED structure, including a GaP window/contact layer, GaInP/InAlP active layers, a Bragg diffraction reflection (DBR) layer, and a GaAs contact layer. An Al_0.95_Ga_0.05_As based sacrificial layer is between the active device and the GaAs substrate for subsequent etching to create freestanding thin-film devices. (c) Circuit model of the optoelectronic resistive sensing based on PR effects. (d) Measured spectra of the excitation source (green light) and PL emissions of the micro-LED connected with different resistors, showing increased PL intensities with resistance (from top to bottom: open-circuit, 650 kΩ, 280 kΩ, 145 kΩ and 120 kΩ). (e) Calculated (lines) and the measured (dots) PL intensity as a function of the load resistance under excitation light with different power densities (3.15 mW/mm^2^ for blue, 1.91 mW/mm^2^ for red and 1.29 mW/mm^2^ for black lines and dots).

**Figure 3** demonstrates the optically measured GSR results of a human subject. The GaInP micro-LED is connected between two different fingers using hydrogel-based wet electrodes (Fig. 3a and more details are in the Figure S3). When a phasic activity (e.g., an action of deep breath) is performed, the skin resistance drops in response to the stimulus,^[23]^ leading to decreased PL intensity compared to the basal condition (Figure 3b). Real-time optical signals are collected in a certain time frame (up to 1000 s), in direct comparison with electrically measured GSR results simultaneously recorded based on a commercial circuit module (Figure 3c and supplement movies S1 and S2). Optical and electrical measurements exhibit an excellent quantitative agreement, showing a determination coefficient of 98.82% when fitting with the theoretical predictions (Fig. 3d). Furthermore, the normalized variation of photon signals (Δ*I*/*I*_0_) is about 4 times larger than electrically measured GSR, demonstrating an inherent capability for amplification associated with the photon-recycling mechanism in the diode. The fluctuation of the measured PL signals is pertinent to various noise sources, which determine the device’s sensitivity. In Figure 3e, the temporal variation of the micro-LED’s PL intensity is recorded when connecting to a constant resistance of 200 kΩ. The noise power spectra lie close to the noise limit of the CMOS camera (Figure 3f), suggesting that the micro-LED introduces negligible instability to the optical sensing system. In accordance with the equivalence relation as shown in Figure 2e, a resolution of 1.1 kΩ Hz^-1/2^ is obtained under this circumstance, by converting the dominant shot noise to the resistance noise. Furthermore, it should be noted that the optical noise is also dependent on the excitation power, which can be further optimized to the trade-off between the signal-to-noise ratio and the GSR sensitivity.

**Figure 3.**
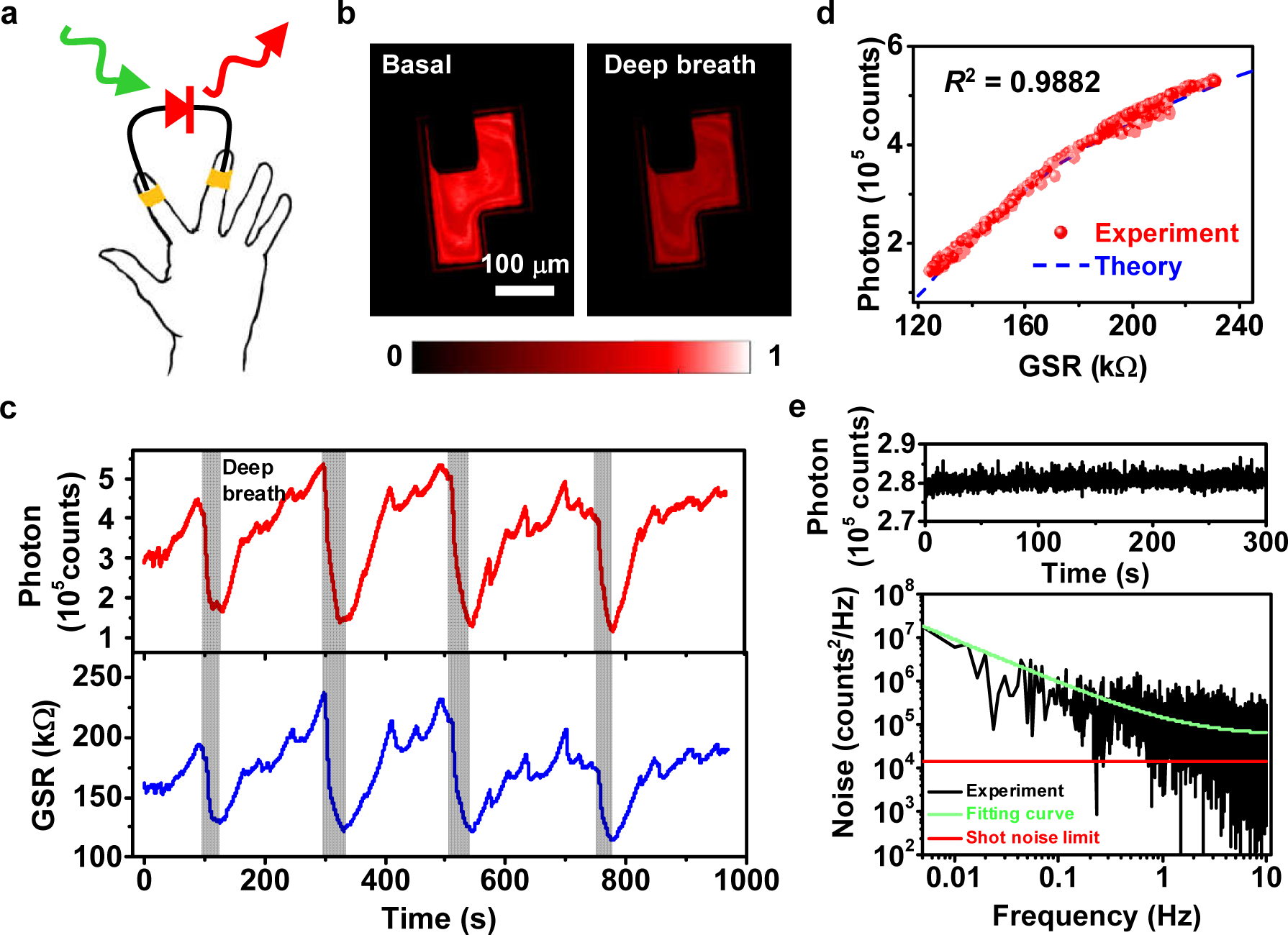
(a) Schematic illustration of the optoelectronic sensing of the galvanic skin response (GSR). (b) Microscopic images of the micro-LED’s PL emission, showing different PL intensities under basal and deep breath conditions of a subject. (c) Simultaneously measured optical signals (top, red) and the electronic signals captured by a commercial GSR sensor (bottom, blue), as a function of time. Gray regions represent discrete deep breath actions of the tested subject, resulting in lower resistance of the epidermis. (d) Measured PL intensity versus GSR, in comparison with the fitting model. (e) The photon counts and (f) the corresponding power spectral density of the PL signals for the micro-LED connected with a 200 kΩ resistor under an excitation power of 1.91 mW/mm^2^.

To summarize, here we demonstrate that electrodermal activities, GSR in particular, can be optically monitored in a circuit-free manner, simply by utilizing a micro-LED. The high photon-to-electron and electron-to-photon conversion efficiencies of III–V semiconductors enable the device to harness the power and emit sensing signals purely optically. In addition, the photon-recycling process in the diode junction endows the device to optically respond to the GSR variation with a high sensitivity. It is envisioned that such a device approach can be further employed to detect many other biophysical and biochemical signals *in vitro* and *in vivo*, with high sensitivity and specificity. Future directions would also involve the formation of devices arrays in the micro- and nanoscale, to realize biosignal mapping at high spatial and temporal resolution. Collectively, we believe that this optoelectronic strategy can offer a novel paradigm for next-generation biosensors in general.

## References

1. D.-H. Kim; N. Lu; R. Ma; Y.-S. Kim; R.-H. Kim; S. Wang; J. Wu; S. M. Won; H. Tao; A. Islam; K. J. Yu; T.-i. Kim; R. Chowdhury; M. Ying; L. Xu; M. Li; H.-J. Chung; H. Keum; M. McCormick; P. Liu; Y.-W. Zhang; F. G. Omenetto; Y. Huang; T. Coleman; J. A. Rogers, Epidermal Electronics. Science 2011, 333 (6044), 838–843.

2. W. Gao; S. Emaminejad; H. Y. Y. Nyein; S. Challa; K. Chen; A. Peck; H. M. Fahad; H. Ota; H. Shiraki; D. Kiriya; D.-H. Lien; G. A. Brooks; R. W. Davis; A. Javey, Fully integrated wearable sensor arrays for multiplexed in situ perspiration analysis. Nature 2016, 529 (7587), 509–514.

3. S. Imani; A. J. Bandodkar; A. M. V. Mohan; R. Kumar; S. Yu; J. Wang; P. P. Mercier, A wearable chemical–electrophysiological hybrid biosensing system for real-time health and fitness monitoring. Nat. Commun. 2016, 7 (1), 11650.

4. J. Kim; A. S. Campbell; B. E.-F. de Ávila; J. Wang, Wearable biosensors for healthcare monitoring. Nat. Biotechnol. 2019, 37 (4), 389–406.

5. H. U. Chung; B. H. Kim; J. Y. Lee; J. Lee; Z. Xie; E. M. Ibler; K. Lee; A. Banks; J. Y. Jeong; J. Kim; C. Ogle; D. Grande; Y. Yu; H. Jang; P. Assem; D. Ryu; J. W. Kwak; M. Namkoong; J. B. Park; Y. Lee; D. H. Kim; A. Ryu; J. Jeong; K. You; B. Ji; Z. Liu; Q. Huo; X. Feng; Y. Deng; Y. Xu; K.-I. Jang; J. Kim; Y. Zhang; R. Ghaffari; C. M. Rand; M. Schau; A. Hamvas; D. E. Weese-Mayer; Y. Huang; S. M. Lee; C. H. Lee; N. R. Shanbhag; A. S. Paller; S. Xu; J. A. Rogers, Binodal, wireless epidermal electronic systems with in-sensor analytics for neonatal intensive care. Science 2019, 363 (6430), eaau0780.

6. M. F. Reynolds; M. H. D. Guimarães; H. Gao; K. Kang; A. J. Cortese; D. C. Ralph; J. Park; P. L. McEuen, MoS_2_ pixel arrays for real-time photoluminescence imaging of redox molecules. Sci. Adv. 2019, 5 (11), eaat9476.

7. D. K. Piech; B. C. Johnson; K. Shen; M. M. Ghanbari; K. Y. Li; R. M. Neely; J. E. Kay; J. M. Carmena; M. M. Maharbiz; R. Muller, A wireless millimetre-scale implantable neural stimulator with ultrasonically powered bidirectional communication. Nat. Biomed. Eng. 2020, 4 (2), 207–222.

8. M. Bariya; H. Y. Y. Nyein; A. Javey, Wearable sweat sensors. Nat. Electron. 2018, 1 (3), 160–171.

9. A. J. Cortese; C. L. Smart; T. Wang; M. F. Reynolds; S. L. Norris; Y. Ji; S. Lee; A. Mok; C. Wu; F. Xia; N. I. Ellis; A. C. Molnar; C. Xu; P. L. McEuen, Microscopic sensors using optical wireless integrated circuits. Proc. Natl. Acad. Sci. U.S.A. 2020, 117 (17), 9173–9179.

10. Z. Lou; L. Wang; K. Jiang; Z. Wei; G. Shen, Reviews of wearable healthcare systems: Materials, devices and system integration. Mater. Sci. Eng. R. Rep. 2020, 140, 100523.

11. D. A. Neamen, Semiconductor physics and devices. McGraw-Hill: New York, 2003.

12. H. Ding; H. Hong; D. Cheng; Z. Shi; K. Liu; X. Sheng, Power- and Spectral-Dependent Photon-Recycling Effects in a Double-Junction Gallium Arsenide Photodiode. ACS Photonics 2019, 6 (1), 59–65.

13. X. Sheng; M. H. Yun; C. Zhang; A. a. M. Al-Okaily; M. Masouraki; L. Shen; S. Wang; W. L. Wilson; J. Y. Kim; P. Ferreira; X. Li; E. Yablonovitch; J. A. Rogers, Device Architectures for Enhanced Photon Recycling in Thin-Film Multijunction Solar Cells. Adv. Energy Mater. 2015, 5 (1), 1400919.

14. M. Peng; Z. Li; C. Liu; Q. Zheng; X. Shi; M. Song; Y. Zhang; S. Du; J. Zhai; Z. L. Wang, High-Resolution Dynamic Pressure Sensor Array Based on Piezo-phototronic Effect Tuned Photoluminescence Imaging. ACS Nano 2015, 9 (3), 3143–3150.

15. M. Stolterfoht; V. M. Le Corre; M. Feuerstein; P. Caprioglio; L. J. A. Koster; D. Neher, Voltage-Dependent Photoluminescence and How It Correlates with the Fill Factor and Open-Circuit Voltage in Perovskite Solar Cells. ACS Energy Lett. 2019, 4 (12), 2887–2892.

16. M. Macka; T. Piasecki; P. K. Dasgupta, Light-Emitting Diodes for Analytical Chemistry. Annu. Rev. Anal. Chem. 2014, 7 (1), 183–207.

17. H. Ding; L. Lu; Z. Shi; D. Wang; L. Li; X. Li; Y. Ren; C. Liu; D. Cheng; H. Kim; N. C. Giebink; X. Wang; L. Yin; L. Zhao; M. Luo; X. Sheng, Microscale optoelectronic infrared-to-visible upconversion devices and their use as injectable light sources. Proc. Natl. Acad. Sci. U.S.A. 2018, 115 (26), 6632–6637.

18. 18. A. Martí; J. L. Balenzategui; R. F. Reyna, Photon recycling and Shockley’s diode equation. J. Appl. Phys. 1997, 82 (8), 4067–4075.

19. O. D. Miller; E. Yablonovitch; S. R. Kurtz, Strong Internal and External Luminescence as Solar Cells Approach the Shockley–Queisser Limit. IEEE J. Photovolt. 2012, 2 (3), 303–311.

20. D. M. Tex; M. Imaizumi; H. Akiyama; Y. Kanemitsu, Internal luminescence efficiencies in InGaP/GaAs/Ge triple-junction solar cells evaluated from photoluminescence through optical coupling between subcells. Sci. Rep. 2016, 6 (1), 38297.

21. C. P. Richter, Physiological factors involved in the electrical resistance of the skin. Am.J.Physiol. 1929, 88 (4), 596–615.

22. W. Boucsein, Electrodermal activity. Springer Science & Business Media: 2012.

23. N. H. Frijda, The emotions. Cambridge University Press: 1986.

